# Drug-Prot: A query system for statistical inference of drug effects and interactions in dynamic proteomic networks

**DOI:** 10.64898/2026.06.17.732914

**Authors:** Markus Ulmer, Rui Sun, Liujia Qian, Ruedi Aebersold, Tiannan Guo, Peter Bühlmann

## Abstract

Understanding drug effects and drug–drug interactions is essential for developing combination therapies. We present `Drug-Prot`, a computational framework that leverages large-scale perturbation proteomics to quantify causal drug effects, drug–drug interactions, and dynamic protein relationships. Using data from 63 single drugs and 59 drug combinations applied to 16 triple-negative and 2 non-triple-negative breast cancer cell lines at 6, 24, and 48 hours, `Drug-Prot` estimates drug effects on protein expression and reconstructs directed temporal protein dependency networks.

The publicly available software enables targeted analyses of user-defined protein sets, substantially reducing the multiple-testing burden. Through an interactive web application, users obtain corrected p-values for single-drug and combination effects, directed temporal dependency networks, and downloadable results without requiring access to the underlying proteomic dataset.

As a use case, we apply invariance-regularized Random Forests to these cancer cell lines to identify proteins associated with drug response. Querying these proteins in Drug-Prot reveals drug-specific and interaction effects at the protein-network level, illustrating how the framework links candidate causal protein features to actionable drug combinations.

**Author Summary:** Cancer is rarely treated by a single drug. Because tumors are driven by networks of interacting proteins rather than by one faulty molecule, treatments are often given in combination. But choosing which drugs to combine is difficult. We usually know what each drug does on its own, and much less about what happens when two are given together.

We analyzed a large experiment in which breast cancer cells were treated with 63 drugs individually and 59 drug pairs, and the levels of more than five thousand proteins were measured at three time points afterward. From this, we estimated which drugs change which proteins, which drug pairs act on proteins in ways that neither drug does alone, and how changes in one protein are followed by changes in another over time.

We have made all of this evidence available through a web tool called Drug-Prot. A researcher can enter a list of proteins they care about and immediately see which drugs and drug combinations affect them, with the statistical uncertainty properly accounted for. We hope this helps others generate and test hypotheses about how drugs affect the proteome.

## 1 Introduction

Drug discovery is an intricate endeavor, largely due to the complexity of biological systems and the multifaceted interactions among drugs, their targets, and downstream molecular networks. Current drug discovery methodologies primarily focus on identifying interactions between drugs and their direct protein targets [1]. However, clinical experience has repeatedly shown that complex and systemic diseases, such as cancer, are rarely controlled through single-target interventions alone. Drug perturbations propagate through protein networks, eliciting adaptive responses that are functionally relevant for therapeutic outcome. Consequently, strategic drug combinations, rather than monotherapy, represent a central approach to addressing the multifactorial nature of complex diseases. Although drug combinations are often pursued for synergistic effects, their clinical relevance remains debatable. Most FDA-approved cancer combination therapies operate predominantly through independent action rather than explicit synergy [2, 3]. Under independent action, drugs do not interact with each other’s mechanisms, and each patient benefits from the drug that works best on their tumor. Our analysis and open-access software enable statistical testing of the additivity and interaction of drug effects on protein expression (as described in more detail below) and thus provide a powerful resource for future studies and validation.

We propose that explicitly modeling the causal interplay among drugs, drug–drug interactions, and protein network dynamics using perturbed proteomic profiles can provide valuable insights into drug discovery and protein-level regulation. Characterizing causal effects is essential, as only causal relationships correspond to the interventional changes induced by a novel drug [4]. Although causal effects are generally not identifiable from purely observational data without strong assumptions [5], the perturbational and temporal structure of the dataset analyzed here enables causal inference under substantially more realistic conditions. Further details are provided in Section 4 (Causal modeling). This framework has the potential to improve the prediction of drug interventions and to identify compatible, non-interfering drug combinations directly from proteomics data by capturing the broader systemic impact of drug perturbations.

The implementation of such frameworks necessitates comprehensive perturbation data that capture functional molecular responses to systematic interventions such as drug treatments or genetic knockouts [6]. However, currently omics datasets available for drug discovery are primarily derived from unperturbed cell models [7–14]. The Connectivity Map (CMap) project represented a major advance in filling this gap by pioneering the systematic profiling of transcriptional responses to drug perturbations [15], subsequently developing the L1000 platform to measure approximately 1’000 landmark transcripts at reduced cost [16]. The Library of Integrated Network-based Cellular Signatures (LINCS) [17] program expects to establish an integrated network-based cellular response database by analyzing changes in gene expression levels and other cellular response levels under a series of perturbations, including transcriptomics [15, 16], proteomics [18, 19], imaging [20, 21], cell viability, and epigenomics data.

While these transcriptome-centric resources have proven valuable, proteins are the primary executors of cellular functions and the direct targets of most therapeutic agents, and early perturbation proteomics efforts have demonstrated their critical role in elucidating drug mechanisms of action (MOAs). However, proteomic measurements within these resources have been limited to only a few hundred proteins [18, 19], leaving essential insights into perturbed protein networks largely unexplored. Ruprecht et al. [22] analyzed the proteomes of 50 drug perturbagens in five lung cancer cell lines after 24 hours, revealing protein variability as a key factor in understanding drug MOAs, particularly with respect to protein homeostasis. Mitchell et al. [23] investigated the perturbed proteomes of 875 molecules in the HCT116 cell line to uncover drug MOAs. Additionally, Zecha et al. [24] provided an approach called decryptM to assay time- and dose-dependent responses of protein and posttranslational modification (PTM) levels after treatment with 31 drugs. Recently, the same group also characterized the dose-resolved perturbation proteomes of 144 drugs in T cells [25]. However, these studies were limited to either single cell lines, single post-treatment time points, or single drug treatments without drug combinations. Recent technological advancements in high-throughput mass spectrometry-based proteomics [26, 27] have enabled profiling of perturbation proteomes across a wider range of paired conditions, including multiple cell lines, various single- and combination-drug treatments, multiple time points, and biological replicates. Leveraging these capabilities, Sun et al. [28] generated a large-scale perturbation proteomics dataset in breast cancer (BCa), encompassing 16 triple-negative and 2 non-triple-negative BCa cell lines, 122 treatments (63 single drugs and 59 drug combinations), measured at 6, 24, and 48 hours after drug treatment, with three biological replicates for each condition.

A small but growing set of platforms exposes drug-perturbed molecular response data through interactive query interfaces (Table 1). The Connectivity Map and its L1000 platform [15, 16] pioneered signature-based querying but operate at the transcriptomic level. On the proteomic side, the Cancer Perturbed Proteomics Atlas [19] couples broad cell-line coverage with the reduced protein coverage of approximately 200 RPPA markers. The LINCS phosphoproteomic and chromatin libraries queryable through Touchstone-P [18, 29] cover only a few hundred analytes. Mass spectrometry signature libraries such as ProTargetMiner [30] achieve near-complete proteome coverage but are restricted to a limited number of cell lines and single time points. The dose- and time-resolved decryptM resource [24, 25] extends the temporal axis but does not include drug combinations. Drug-combination synergy databases such as DrugComb [31] and DrugCombDB [32] address combination evidence at the viability-readout level and are complementary to, rather than directly comparable with, the proteomic platforms summarized in Table 1.

**Table 1:** Comparison of `Drug-Prot` with related platforms that expose drug-perturbed molecular response data through interactive interfaces. “Yes” indicates the platform supports the feature as a first-class capability, “partial” indicates limited support (e.g. reduced sub-proteome, narrow scope, or evidence not statistically calibrated), “limited” indicates the underlying data contain the feature, but the platform exposes it only indirectly, and “no” indicates that the feature is absent.

| **Feature** | **Drug-Prot**<br>(this work) | **CPPA**<br>[19] | **TouchstoneP**<br>[18, 29] | **ProTarget Miner**<br>[30] | **decryptM**<br>[24, 25] | **CMap / L1000**<br>[15, 16] |
|---|---|---|---|---|---|---|
| Proteomic readout | yes | partial | partial | yes | yes | no |
| Near-complete proteome coverage | yes | no | no | yes | yes | no |
| Multiple cell lines | yes | yes | yes | limited | yes | yes |
| Multiple post-treatment times | yes | limited | limited | no | yes | limited |
| Drug combinations in data | yes | limited | no | no | limited | no |
| Statistical inference exposed | yes | yes | no | no | partial | no |
| Drug–drug interaction testing | yes | no | no | no | no | no |
| Directed protein-dependency network | yes | no | no | no | no | no |
| Interactive query on user-defined protein set | yes | partial | partial | partial | no | partial |

`Drug-Prot` differs from these resources in two respects. It provides calibrated p-values for drug–drug interaction effects on user-defined protein sets and exposes directed temporal protein-dependency networks inferred from drug perturbations across multiple cell lines. Connectivity-based metrics, such as the *τ* -score used in the CMap rank query, reference signature similarity against an empirical background of observed signatures. `Drug-Prot`, by contrast, reports calibrated p-values for the null hypothesis of no effect, with explicit multiple-testing correction applied at the level of the user-defined protein set. To our knowledge, none of the existing platforms support protein-set-level queries with corrected statistical inference, a feature particularly valuable for pathway-focused and hypothesis-driven analyses.

### 1.1 Drug-Prot

Leveraging the systematically designed perturbation proteomics dataset from Sun et al. [28], `Drug-Prot` is a computational framework that statistically quantifies the causal effects of single-drug perturbations and drug pairs on protein expression, and complements them with a directed temporal dependency network between proteins, inferred from the dynamics of the perturbation experiments across multiple time points. While drug effects can be interpreted causally under the experimental intervention design, the inferred protein dependencies are directional and temporally ordered associations that may still reflect unmeasured biological confounding. Consequently, these networks should be viewed as hypothesis-generating rather than mechanistic. The diversity of interventional perturbations, coupled with the temporal nature of the proteomics measurements in this dataset, enables causal-type modeling of various effects.

A key novelty of our method lies in its ability to incorporate user-defined protein sets as the focus of the analysis. That is, one can query the effects of drugs for some proteins of interest, and `Drug-Prot` outputs the desired significance results. By allowing the researcher to specify proteins of interest, `Drug-Prot` produces tailored outputs that highlight significant drug effects on the selected proteins, both from individual drugs and from drug combinations. Furthermore, the framework captures dynamic aspects of protein behavior by identifying directed temporal dependencies between proteins that evolve from one time point to the next. This dual perspective, linking drug perturbations to changes in temporal interactions, provides a powerful tool for studying the molecular mechanisms of drug action in a targeted yet comprehensive manner.

The underlying models are fitted once to all 5’078 proteins, and the resulting p-values serve two uses. Corrected across the whole proteome, they give an overview of the dataset, which we summarize in the Results. Corrected across a protein set selected by the user, they give a focused analysis with more power, because the multiple-testing correction covers only the hypotheses of interest.

## 2 Results

### Overview of

`Drug-Prot`. This section provides an overview of `Drug-Prot` while the Methods section describes the underlying model and methodology in detail. In particular, when we use the term ‘causal effects’, whether direct or indirect via mediation through other quantities (total effect), this necessarily must be with respect to certain models.

Drug-Prot is a computational framework equipped with an interactive Shiny application. Its output consists of two primary components. First, it estimates the direct and total causal effects of single drugs and drug interactions on proteins of interest. Specifically, the framework reports statistical p-values for differential protein expression (relative to untreated controls) at 6, 24, and 48 hours following drug administration. Second, Drug-Prot quantifies temporally ordered, directed dependencies between proteins, providing p-values for differential expression attributable to protein activity at earlier time points. When the user supplies a protein set, the dependency analysis is not restricted to relationships within that set. For every queried protein, Drug-Prot considers every protein in the proteome (including the other queried proteins themselves) both as a potential parent at the preceding time point and as a potential child at the subsequent time point. The returned subgraph and p-values therefore describe how each queried protein relates, in either temporal direction, to any other protein measured in the dataset.

A schematic description of the framework is given in Figure 1(a). While the initial training of `Drug-Prot` is based on the proteomic dataset, any use of `Drug-Prot` is based on the information of approximately 55 million p-values for drug and protein effects for all 5’078 proteins that are included in the dataset, without the need to access the initial data. A key feature of `Drug-Prot` is that users can predefine a set of proteins of interest, provided they are present in the dataset. This functionality not only enhances the interpretability of the results but also substantially mitigates the multiple-hypothesis testing problem, thereby reducing the need for overly conservative p-value corrections. Accessibility of `Drug-Prot` is described in Section 5.

**Figure 1:**
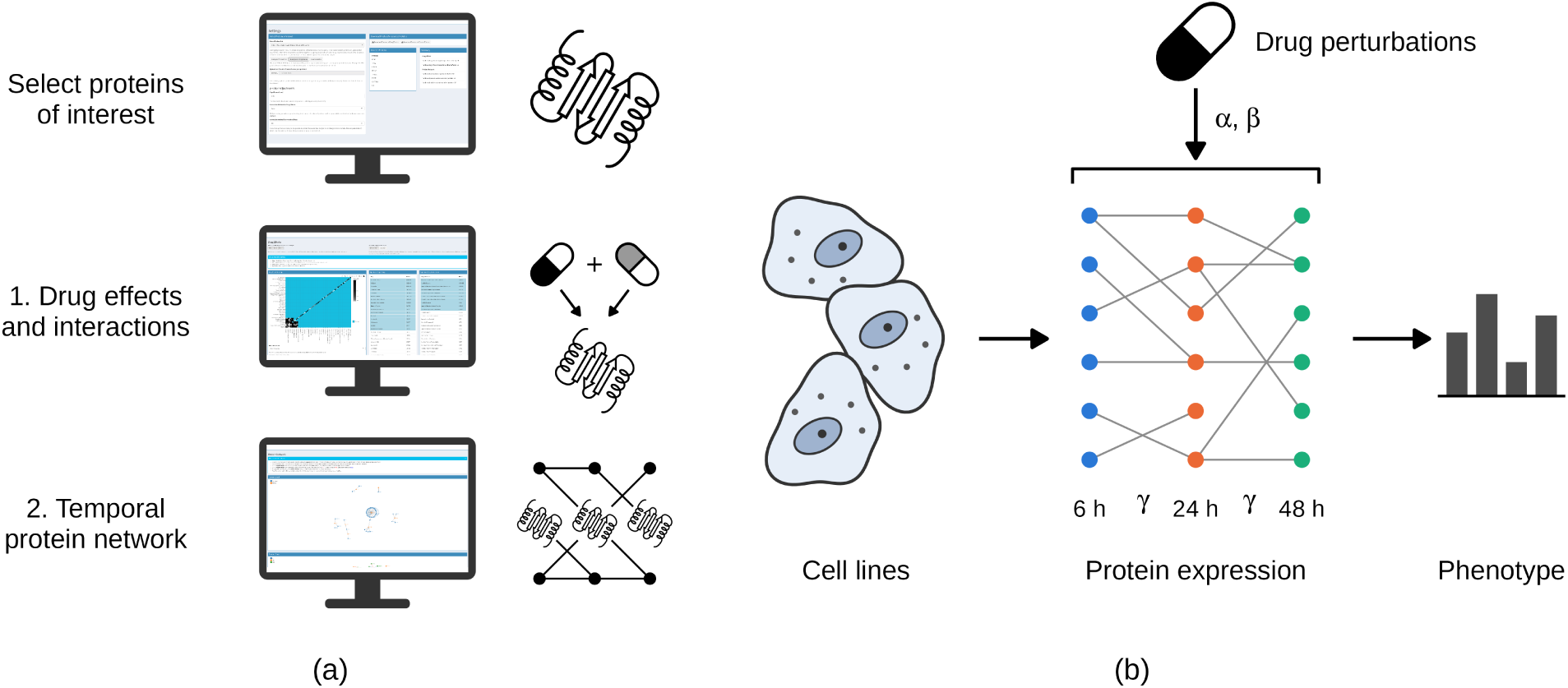
Schematic visualization of Drug-Prot. Figure 1(a) shows the software interface to query proteins of interest. Given a set of proteins, Drug-Prot lets a user choose a significance level and the desired p-value correction. It then indicates, first, which drugs show a significant effect on the selected set of proteins and which drugs interact with respect to these proteins. Second, it presents the directed temporal dependency network for the selected proteins and their connected proteins, in both temporal and summary graphs. In Figure 1(b), a schematic of the available data and assumed models is shown. The assumed model for the drug perturbation effects on the protein expression follows equation (1) and the protein-to-protein temporal dependencies follow equation (2).

Our analysis focuses on the following key quantities:

1. **Causal drug effects on protein expression**. We evaluate protein differential expression relative to baseline at 6, 24, and 48 hours post-treatment and report statistical evidence for causal effects. At 6 hours, these are total effects. At 24 and 48 hours, Drug-Prot reports both the total effect and the direct effect, which excludes the part of the effect mediated by the proteome at the preceding time point. Statistical evidence is reported separately for single-drug interventions and two-drug interactions. Figure 2(a) illustrates the corresponding p-values over all time points for a selected set of proteins. In this visualization, the diagonal entries show the statistical significance of single-drug treatments, while the off-diagonal entries show two-drug interactions. Drug pairs for which we lack data are shown in cyan, outside the p-value color scale. Figure 2(b) shows a zoom-in of Figure 2(a) on the combinations for which we have experimental data. These p-values quantify the significance of causal effects for proteins within a pre-specified set, with appropriate adjustments for multiple testing across proteins and time points. When the set contains only a single protein, the reported p-value pertains directly to that specific protein. The figures and counts in this section use the direct effects at 24 and 48 hours, which separate the action of a drug from the response it propagates through earlier protein changes. The corresponding total effects are available in Drug-Prot.
2. **Directed temporal dependencies in the protein network**. We assess, for every pair of proteins, whether differential expression of one protein at *t* = 6 or *t* = 24 hours is significantly associated with differential expression of the other at the subsequent time point (*t* = 24 or *t* = 48 hours, respectively), after adjusting for the residual drug effects in model (2). A significant association is reported as a directed edge from the earlier to the later (protein, time) node in the dependency graph (Figure 3(b)). We emphasize that the direction of an edge reflects the temporal ordering of the measurements rather than a verified causal mechanism. Hidden confounders acting between time points may still influence both endpoints.

**Figure 2:**
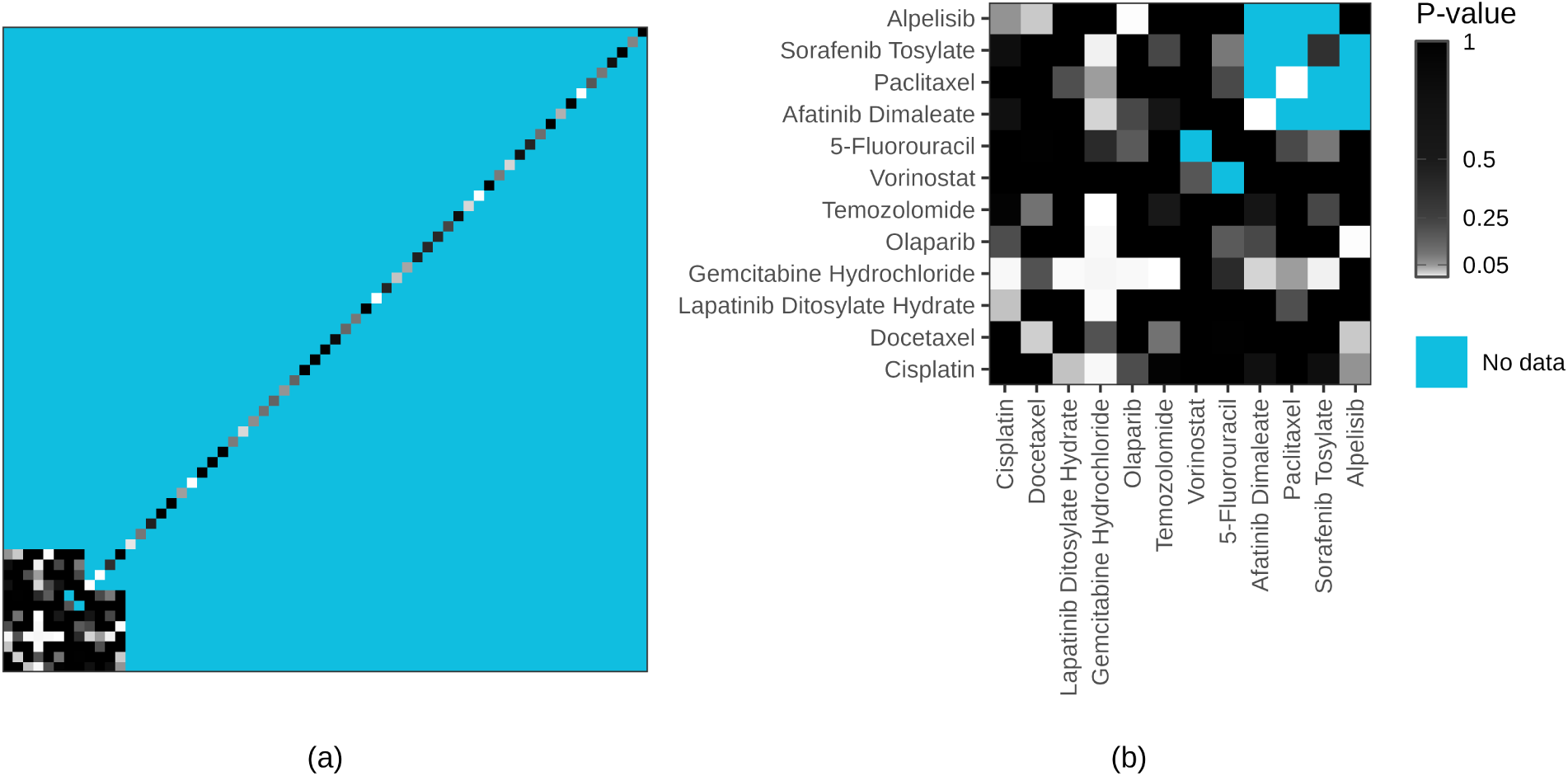
Sixteen single drugs and ten drug pairs act on the eight IC50-predictive proteins, and seven of the ten pairs contain Gemcitabine. The heatmap shows corrected p-values for the drug effects on the eight proteins selected by the Anchor Forest in Section 4.1. The diagonal elements contain the minimal corrected p-value for the corresponding single drug across the three time points and the eight proteins, testing whether at least one protein at at least one time point is affected by the drug. The off-diagonal elements correspond to the overall p-value for a drug interaction. The minimum is taken over the drug effects of model (1) at 6 hours and the direct drug effects of model (2) at 24 and 48 hours. Drug combinations for which we lack experimental data are shown in cyan, and p-values from 0 to 1 are colored from white to black. Figure 2(b) zooms in on the drug combinations for which we have data. The significant single drugs at the 5% level are Entrectinib, Niraparib Tosylate, Afatinib Dimaleate, Bosutinib Monohydrate, Dasatinib Hydrate, Verteporfin, Raloxifene Hydrochloride, Docetaxel, Doxorubicin Hydrochloride, Anastrozole, Gemcitabine Hydrochloride, Irinotecan Hydrochloride, Paclitaxel, Estradiol, Vincristine Sulfate, and Methotrexate.

**Figure 3:**
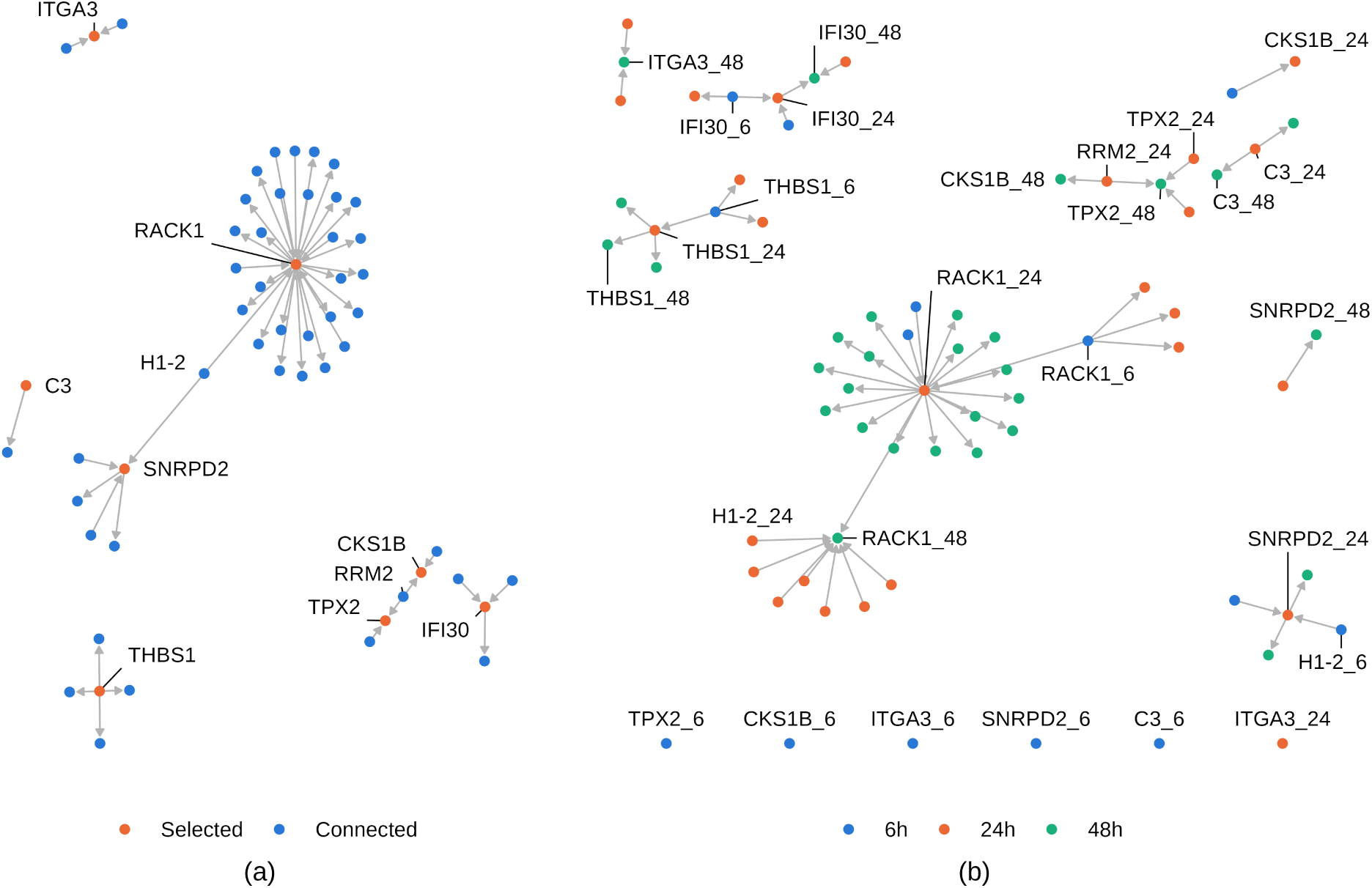
Temporal dependencies around the eight IC50-predictive proteins. Figures 3(a) and 3(b) show the summary and temporal graphs of the directed temporal protein dependencies of model (2) for the eight proteins selected by the Anchor Forest in Section 4.1. In the temporal graph in Figure 3(b), protein expression at 6 hours (blue) is linked to 24 hours (orange), and 24 hours to 48 hours (green). An edge is drawn whenever the p-value of the corresponding *γ* parameter is significant at a false discovery rate of 5%. Edge direction reflects the temporal ordering of the measurements and is not protected against unmeasured confounding between time points. The summary graph in Figure 3(a) draws an edge between two proteins if there is a corresponding edge in the temporal graph. The selected proteins are shown in orange, and the proteins directly connected to them in blue. A label such as TPX2 48 gives the protein and the time point of the measurement in hours. Selected proteins without a significant dependency at a time point are drawn in a row below the graph.

### Statistical evidence in the full dataset

We present a comprehensive analysis of all 5’078 proteins in the dataset. For the key quantity described in item 1 above, we obtained the following results. Across the 122 drug treatments and 5’078 proteins (yielding 1’858’548 potential effects), we identified 5’475 significant (at a family-wise error rate of 0.05) drug effects at 6 hours, 244 at 24 hours, and 294 at 48 hours. Most significant drug effects occur at 6 hours. At 24 and 48 hours, total effects remain frequent, with 7’088 and 4’436 significant effects, whereas direct effects are rare, indicating that later changes in protein expression largely propagate through earlier protein responses rather than through residual drug action. At 6 hours, most significant single-drug effects concentrate on a small group of proteins, which respond in the same direction to most drugs relative to the time-matched reference. The effects at 24 and 48 hours are direct residual effects, adjusted for changes that propagate through earlier protein expression responses. As discussed in Section 4 (Causal modeling), the causal reading of these residual effects at 24 and 48 hours rests on the additional Assumption 2, so they carry more uncertainty than the total effects. Among all interaction effects of two drugs across all proteins (299’602 effects per time point), 0.42% at 6 hours, 0.038% at 24 hours, and 0.042% at 48 hours show a significant interaction term.

For the quantity described in item 2 above, we construct a temporal graph representing directed temporal dependencies between proteins, specifically from 6 to 24 hours or from 24 to 48 hours. The total number of possible directed dependencies is 51’572’168. Among these, we identify 26’092 significant dependencies (at a false discovery rate = 0.05). We summarize these connections in a summary graph, drawing an edge between two proteins if they are significantly linked at any time point. In the summary graph, there are 25’781’006 possible directed connections, of which 23’692 are significant, corresponding to an average network degree of 9.3. These are summary statistics of the output of `Drug-Prot` overall. Examining a smaller set of proteins is often more meaningful, as discussed next.

### A use case with eight pre-selected proteins

We illustrate Drug-Prot with a predefined set of proteins, the eight proteins with the largest variable importance in the invariance-regularized Random Forest for IC50 response, see Section 4.1. The selected proteins are ITGA3 measured at 6 hours, CKS1B at 24 hours, and TPX2, IFI30, THBS1, RACK1, SNRPD2, and C3 at 48 hours. The regularization favors predictions with similar residuals across treatments, targeting proteins with a stable relationship to IC50 under drug perturbations. Invariance under the observed drug shifts is a property of causal relations but does not prove one, so we treat the eight proteins as candidate markers of drug sensitivity. We then use Drug-Prot to infer the effects of drug treatments on these candidates, which points to treatments that may modulate IC50 through them.

When examining key quantity 1, **causal drug effects on protein expression**, we obtain the corresponding p-values for both single and paired-drug treatments (Figure 2(a)). Applying a 5% family-wise error rate threshold, 26 out of 122 possible drug effects are significant at one or more time points (6, 24, or 48 hours post-treatment). Of these, 16 are single-drug effects, and 10 are two-drug interactions, so 49 of the 59 evaluated drug pairs show no evidence of interaction for this set of proteins. The strongest single-drug effects come from Afatinib Dimaleate (*p* = 1.4 × 10^−9^), Paclitaxel (*p* = 4.8 × 10^-6^), Verteporfin (*p* = 5.3 × 10^-5^), Vincristine Sulfate (*p* = 1.9 × 10^-4^), Entrectinib (*p* = 4.0 × 10^-4^), and Dasatinib Hydrate (*p* = 5.2 × 10^-4^).

Gemcitabine Hydrochloride stands out among the interactions. It affects the set on its own (*p* = 0.0022) and shows evidence of interaction with seven of its eleven partners, namely Temozolomide, Lapatinib Ditosylate Hydrate, Olaparib, Cisplatin, Sorafenib Tosylate, Afatinib Dimaleate, and Paclitaxel. Paclitaxel, in contrast, acts on the set with a strong effect but shows evidence of interaction only with Gemcitabine, and its seven other combinations show no evidence of departure from additivity. Gemcitabine is a nucleoside analog that depletes deoxynucleotide pools through inhibition of ribonucleotide reductase and terminates DNA synthesis after incorporation [33]. Its widespread interactions with drugs of different classes are consistent with an action on a shared resource of the cell, the nucleotide supply, which most of its partners also tax, directly through DNA damage or its repair (Cisplatin, Temozolomide, Olaparib) or indirectly through proliferation signaling (Lapatinib, Afatinib, Sorafenib). Two of the seven combinations have also shown efficacy in clinical studies. Gemcitabine combined with Paclitaxel prolongs survival relative to Paclitaxel alone in metastatic breast cancer after anthracycline treatment [34], and Gemcitabine combined with Cisplatin is a first-line option for metastatic triple-negative breast cancer [35].

For key quantity 2, **directed temporal dependencies in the protein network**, we present results in Figure 3(b). At a false discovery rate of 5%, the summary graph in Figure 3(a) for the eight pre-selected proteins reveals 48 significant connections out of 81’176 possible, corresponding to an average degree of 6, and the temporal graph in Figure 3(b) identifies 56 significant connections out of 162’368. THBS1, RACK1, C3, TPX2, and IFI30 depend on their own earlier expression, which reflects the persistence of expression changes over time. Beyond these self-dependencies, RRM2 measured at 24 hours is an antecedent of both TPX2 and CKS1B at 48 hours, with positive coefficients (*p* = 4.4 × 10^−4^ and *p* = 0.009). RACK1 is the most connected protein of the set, with positive edges to proteins of replication, protein folding, and proteasomal degradation, such as PCNA, CCT5, PSMD14, and HDAC1. While these results are of a statistical nature, their biological plausibility is discussed in S1 Appendix, Section B. As a check against known pharmacology, S1 Appendix, Section C queries nine drug targets in 21 drug-target pairs with a predicted outcome, and Drug-Prot recovers five of the twelve predicted changes with the expected direction and time course and reports a change for none of the nine pairs predicted to stay unchanged at 24 or 48 hours.

## 3 Discussion

Based on the large-scale perturbation proteomics dataset generated by Sun et al. [28], `Drug-Prot` provides statistical evidence for drug effects and drug–drug interactions on protein expression (interpretable as causal under externally controlled drug administration) together with a directed temporal dependency network between proteins, interpretable as directional but potentially confounded. All statistically inferred drug–protein and protein–protein relationships covered by the data are accessible via `Drug-Prot`. This enables users to explore, validate, and reinterpret results beyond those explicitly discussed in the text.

We present a use case with eight selected proteins that are predictive of the IC50 phenotype across drug perturbations and thus candidates for causal effects on it. We also provide statistical validation of the findings and a comparison with known drug effects on target abundance (S1 Appendix, Section C), which supports the potential of Drug-Prot to yield reasonable biological insights. Any other mechanism or prior knowledge can be used to select a set of proteins of interest. In this sense, Drug-Prot serves as a mechanism for disseminating a substantially richer body of statistical evidence than can be reasonably presented in a single publication. Users can interrogate specific proteins, drugs, or drug combinations of interest, examine associated interaction structures, and assess the significance of hypotheses about causal effects, provided the underlying dataset from [28] covers them.

### Direct and total drug effects

`Drug-Prot` reports the significance of drug effects at 6, 24, and 48 hours. At 24 and 48 hours, each effect is available as a total effect and a direct effect, and the choice depends on the research question. At 6 hours, the two coincide. The total effect describes the full change in a protein’s expression after treatment, including changes that propagate through earlier protein responses. It answers pharmacological questions, such as whether a drug alters a candidate biomarker, and its causal reading needs only Assumption 1 (Section 4). The direct effect describes the part of this change that is not mediated by the proteome measured at the preceding time point. It is closer to a mechanistic question, such as whether a drug acts on a protein beyond the cascade it triggers earlier, but it relies on the stronger Assumption 2 (Section 4). A significant total effect without a significant direct effect at a later time point therefore points to an effect propagated through earlier protein responses, although non-rejection can also reflect limited power.

### Use cases

While we present a single use case in detail (the IC50 phenotype in Section 2), `Drug-Prot` is intended as a general resource for investigating biologically motivated protein sets. The examples below illustrate a range of queries that can be addressed directly through the platform, highlighting its general utility.

- *DNA damage response.* A user studying genotoxic agents can query a curated DNA-damage-response set and ask which single drugs and which drug pairs significantly perturb it, and how its members are linked across time.
- *Drug-sensitivity markers.* Proteins associated with drug sensitivity, such as the eight IC50-predictive proteins studied in Section 2, which were derived from 16 triple-negative and 2 non-triple-negative cell lines, can be analyzed for the treatments most likely to modulate response and for the drug pairs whose effect departs from additivity on them.
- *Combination partners of a drug.* Starting from a drug of interest, a user can ask which partners show evidence of interaction on a protein set, as for Gemcitabine in Section 2, where seven of eleven partners do.
- *Autophagy and endolysosomal pathways.* A pathway-level set, for example the proteins of endolysosomal trafficking, can be uploaded as a plain-text list to obtain corrected evidence at the pathway scale, where proteome-wide corrections would be prohibitively conservative.

Any pathway, protein complex, or curated gene set (e.g. from KEGG, Reactome, or MSigDB) can be queried in the same way. Because the multiple-testing correction is applied only over the queried set, such focused analyses gain substantial power relative to proteome-wide searches.

Beyond specific queries, the directed temporal dependency network derived from Drug-Prot enables systematic exploration of indirect relationships and potential mediating pathways. This network structure supports downstream analyses using graph-based methods, including community detection, pathway-level aggregation, and flow-based approaches, allowing users to contextualize statistical findings within broader molecular architectures.

### Outlook

The time points of 6, 24, and 48 hours are those of the dataset of Sun et al. [28], and the models are not tied to them. Model (1) applies at every measured time point and model (2) applies to every pair of consecutive time points. The same analysis could therefore be applied to other perturbation datasets with externally assigned treatments, untreated controls, and measurements at two or more time points.

One such case would be patient-derived models with known clinical outcomes. There, protein sets whose drug response differs between responders and non-responders could be tested as markers of response to therapy, in analogy to the eight proteins of Section 2, which predict IC50 in culture. The present data cannot support this step. It comes from 18 established cell lines in culture, not patients, and the response measure is IC50 in culture, not response to therapy. In addition, three biological replicates per condition are insufficient to estimate drug effects for each cell line separately.

## 4 Methods and Statistical Validation

Underneath `Drug-Prot` are p-values for parameters associated with drug effects and protein expressions at the most recent time points. These p-values are obtained from high-dimensional linear models. In each case, the response variable corresponds to a single differential protein expression, and this procedure is repeated for all 5’078 measured proteins.

### Structure of the dataset and notation

We measure expressions of 5’078 proteins at different time points *t* ∈ {0, 6, 24, 48} (in hours). The protein expressions for time point *t* are denoted by *P^t^* ∈ ℝ*^nt^*^×5078^. We distinguish between *t* = 0, the baseline case with no drug perturbation, and *t >* 0, where drug perturbations were administered, and protein expression was measured after *t* hours. The samples are from *k* ∈ {1*, . . . ,* 18} cell lines, and when *t* ≠ 0, drug perturbations were applied with certain drug concentrations *D* ∈ ℝ*^nt^*^×63^. We have 4’625 samples at *t* = 6, 4’649 at *t* = 24, and 4’540 at *t* = 48. We denote a sample point by *i* ∈ {1*, . . . , n_t_*}, and we sometimes call it an experiment: it corresponds to a cell line, a drug perturbation, and a specific replicate for that cell line and perturbation. A sample is observed only at a single time point due to the destructive nature of protein measurement, and we thus have unpaired data across different time points *t*. Therefore, we consider a sample point or experiment *i* only for a single and fixed time point *t*, and we denote by *D_i_* ∈ ℝ^63^ the drug concentration vector for the sample *i* and by *C_i_* ∈ {1*, . . . ,* 18} the index of the cell line.

If a single drug was administered for sample *i*, there is only one non-zero element in the *i*-th row of *D*, and it always has the concentration being equal to 10 µM: that is, for a single element *j* ∈ *J*, where *J* = {1*, . . . ,* 63}, we have that *D_i,j_* = 10 and all other entries are zero, with *D_i,j_′* = 0 for all *j* ≠ *j*. The other perturbation samples or experiments were administered with two drugs, that is, a drug combination, whose concentrations are chosen between 0.00025 and 30 µM, and thus, for such a sample *i*, exactly two entries of the *i*-th row of *D* are non-zero.

As mentioned above, the data is unpaired over different time points. Thus, whenever a protein expression is modeled using protein expressions at other (previous) time points, we must use an aggregated version of the latter. Consider a protein *r* ∈ {1*, . . . ,* 5078}. We denote the aggregated version of a log-protein expression given an experimental condition *C_i_, D_i_* in sample *i* by 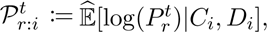 where Ê stands for the sample median within the corresponding experimental condition. We use the median to be robust to various unwanted sources of variation, such as batch effects or measurement errors.

A differential protein expression 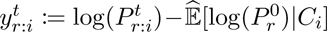 at any time point *t >* 0 is always defined relative to its baseline expression level prior to treatment. We denote the aggregated version of the differential log-protein expression by 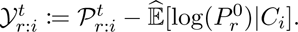

A fuller description of the dataset, its design, scale, and acquisition, is given in the S1 Appendix, Section A.

### Drug effects on differential protein expression (6 hours post-treatment)

Here, we model differential expression levels at 6 hours as a function of drug administration

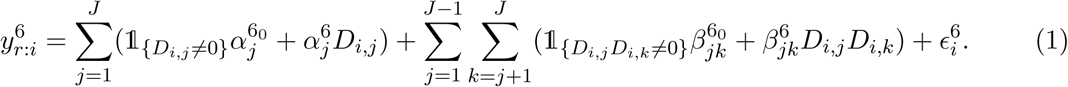

For treatments administered at a single concentration, the intercept and the concentration term are not separately identified, and we keep only the intercept. Here the *α* coefficients measure single-drug effects, and the *β* coefficients measure drug-interaction effects. *α*^0^ and *β*^0^ are treatment intercepts, while *α* and *β* are linear coefficients of the drug concentrations. The sum over *j < k* runs over unordered drug pairs, so each pair contributes a single interaction term. Because a single-drug experiment has exactly one non-zero entry in the corresponding row of *D*, the product *D_i,j_D_i,k_* vanishes for all single-drug samples. The interaction parameters are therefore identified only from the experiments in which that pair was co-administered, which is why interaction effects are testable for the 59 measured pairs. These parameters encode departure from additivity on the differential log-expression scale. If both 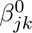 are zero, the effect of the combination on protein *r* is the sum of the two individual effects. A non-zero value indicates that the combined effect departs from this sum. *ɛ* are i.i.d. error terms with E[*ɛ_i_*] = 0. All these quantities depend on *r*, but we suppress this in the notation.

The parameters are estimated using de-sparsified Lasso [36, 37] implemented in the R package *hdi* [38]. Group p-values are computed for both single and interaction effects, as the minimum of the p-values of the parameters of a treatment, multiplied by the number of these parameters (Bonferroni correction).

### Protein expression dynamics across time points (6 to 24 hours, 24 to 48 hours)

Here, we model differential expression levels 24 and 48 hours after treatment as a function of the protein differential expressions at the previous time point and the residual drug effects:

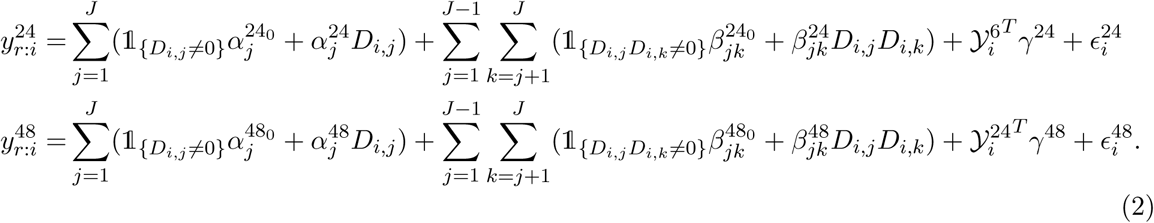

For the definition of *Y*, see the end of the subsection Structure of the dataset and notation. In addition to the drug effects, we now also have the parameters *γ*^24^ and *γ*^48^, which capture the temporal effects of proteins. Meanwhile, *α* and *β* now only capture residual drug effects that are not mediated through proteins at previous times. Group p-values are again computed for both single and interactive drug effects, as well as single p-values for protein expression dynamics, using the same high-dimensional inference framework.

Model (2) estimates the drug effects at 24 and 48 hours conditional on the proteome at the preceding time point. To obtain the drug effects at these time points without this conditioning, we also fit model (1) with the response 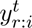 for *t* ∈ {24, 48}. We denote its parameters by *ᾱ^t^*^0^, *ᾱ^t^*, *β̄^t^*^0^ and *β̄^t^*, and compute the group p-values in the same way as at 6 hours.

### Set of proteins of interest and multiple testing

Given a set of proteins of interest `R` of size *M*, we have 122×3×*M* p-values for drug effects (122 single and pairs of drug effects at 3 time points) and *M* × 5078 × 2 + *M* × (5078 − *M*) × 2 p-values for protein on protein effects (the first summand are the p-values for parents of `R` and the second summand are children of `R`). In the case of drug effects, we control for family-wise error rate (FWER) by applying a Bonferroni-Holm correction [39]. For protein-to-protein temporal dependencies, we control the false discovery rate (FDR) using the method of [40]. We use FWER control for drug effects because the number of tests is moderate (e.g., 366 for a single queried protein), so power remains adequate, and FWER control allows us to interpret each significant drug or drug pair individually. For the protein–protein dependencies, the number of tests per query is very large (e.g., 20’310 for a single protein and 162’368 for the eight-protein set), so we treat this as large-scale screening and control the FDR instead. Alternatively, in `Drug-Prot`, users can select any of `R`’s `p.adjust` correction methods (Holm, Hochberg, Hommel, Bonferroni, Benjamini–Hochberg, Benjamini–Yekutieli), or no correction, for downstream tasks.

### Interpretation of p-values and significance

The group p-values for *α* and *β* provide valid statistical tests of whether an effect is non-zero for at least one selected protein and at least one selected time-point. So, if *α* is significant, we have sufficient evidence that the corresponding drug affects at least one of the selected proteins at at least one time point. A significant *β* gives significant evidence of a deviation from the additive model in these cases. A non-significant result does not mean a drug does not affect a set of proteins, and it does not mean two drugs do not interact. Non-rejection can reflect a true absence of effect, but it can equally reflect insufficient power due to low sample size. The same holds for the significance statements about *γ*.

Figure 4 examines the validity of the reported p-values in synthetic data that uses the observed design and estimated parameters from the real data, but with simulated Gaussian noise. We sample 100 response proteins at random and simulate their responses at all three time points from the model with estimated coefficients, using Gaussian noise with estimated residual variances and keeping the design fixed.

**Figure 4:**
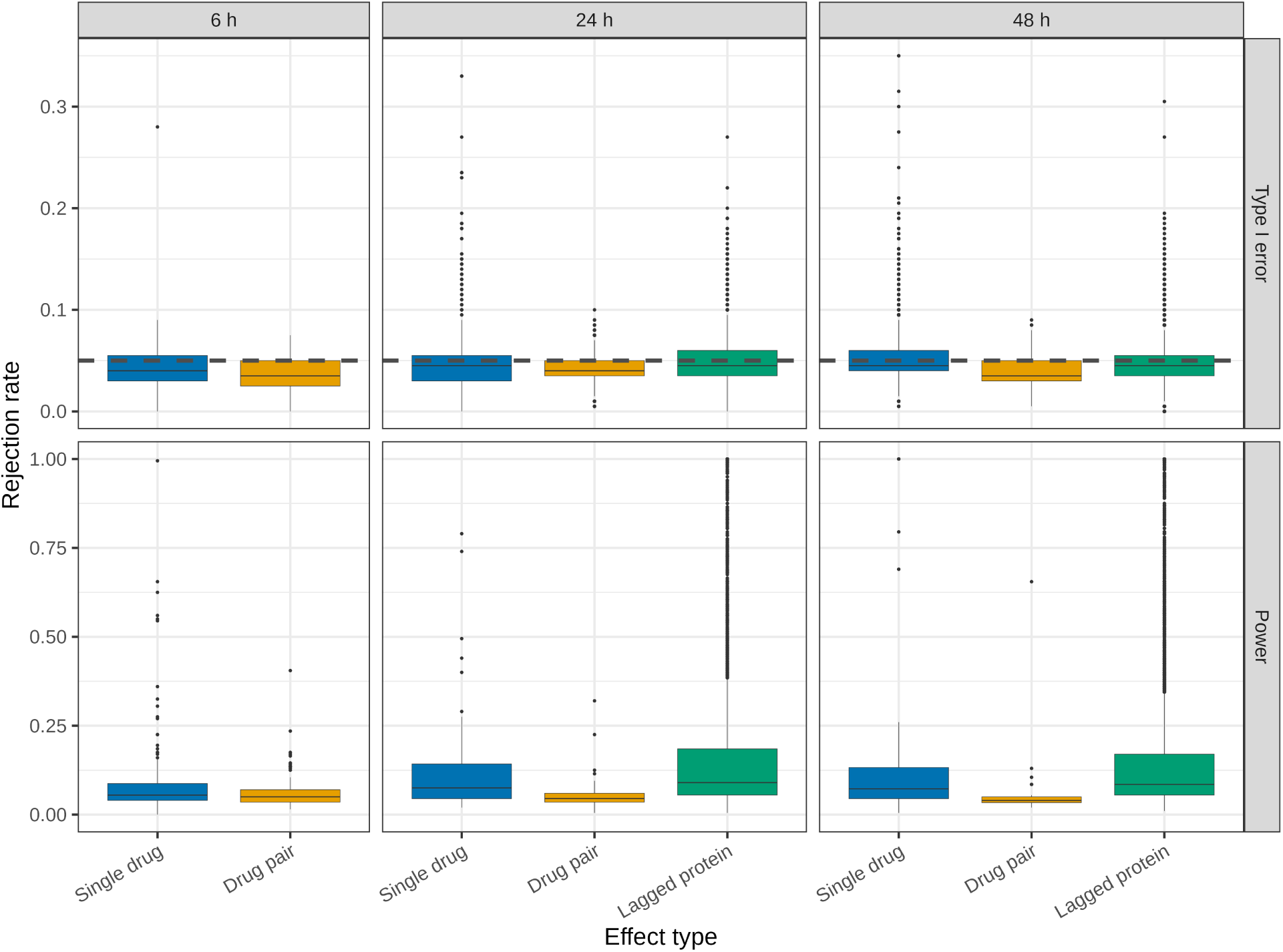
Rejection rates of p-values in synthetic data, sampled from the corresponding models with estimated parameters from real data. Rejection rates at level 0.05 over 200 simulated data sets for 100 randomly chosen response proteins, for coefficients that are zero (type I error, top) and non-zero (power, bottom) in the simulation. The dashed line marks 0.05. Drug tests are slightly conservative, and clearly inflated rates are limited to a few single drugs at 24 and 48 hours and some protein effects.

In each simulated data set, we refit the model and compute the group p-values for the drug effects and the individual p-values for the protein effects as in the original analysis. Over 200 repetitions, the rejection rate at level 0.05 estimates the type I error for coefficients that are zero in the simulation and the power for the others. The drug tests are slightly conservative, with median rejection rates between 0.035 and 0.045 under the null. For the lagged proteins, the median rejection rate under the null is 0.045, and 99% of the predictors stay below 0.085, within the binomial variation from 200 repetitions. Clearly inflated rates occur for a few single drugs in the direct-effect models at 24 and 48 hours. Most of these tests involve covariates whose nodewise residual *Z_j_* remains strongly correlated with another covariate, which Zhang and Zhang [36] quantify with the bias factor 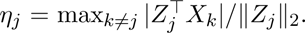 We report *η_j_* for every test in the deposited evidence files, so users can check individual results.

If a user selects proteins, a significance level, and a correction scheme before looking at the results, the results on drug effects and temporal dependencies between proteins provide valid statistical tests. In practice, users will interact much more freely with the platform, and tests chosen after seeing results lose their validity. This also applies to a protein set assembled from the most significant proteome-wide results and extended by known interaction partners. We believe the p-values, including the full set of uncorrected p-values, remain informative for generating and prioritizing hypotheses in this case.

### Causal modeling

The two kinds of parameters in models (1) and (2) carry different interpretations. Drug-effect parameters describe what happens under an intervention. Protein-to-protein parameters describe temporally ordered dependence that may be confounded. Figure 5 shows the assumed structure.

**Figure 5:**
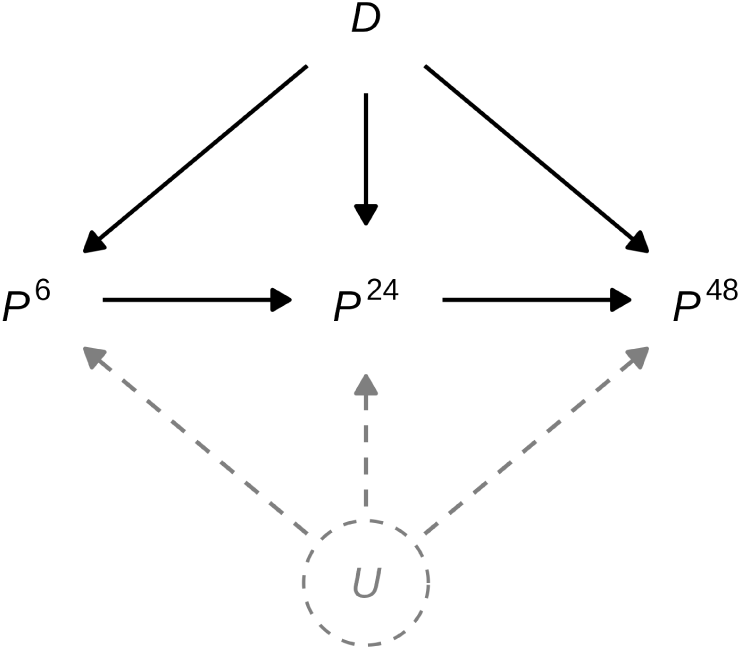
Assumed causal structure in the drug perturbation models. *D* is the administered drug perturbation, a single drug or a drug pair. *P* ^6^, *P* ^24^ and *P* ^48^ are vectors of differential protein expression measured 6, 24 and 48 hours after treatment. Solid edges from *D* to each *P^t^* are direct interventional effects of treatment. Solid edges between successive expression nodes are temporally ordered dependencies. No edge points into *D*, which reflects the assumption that treatment is assigned externally. The dashed node *U* stands for unmeasured biological processes that may influence expression at several time points, such as proteins outside the measured set. *U* is drawn to make explicit what the models do not adjust for.

The experimental design supports an assumption that enables inferring total causal effects, namely the following.

#### Assumption 1

No hidden confounder has a directed edge into the drug treatment *D*, that is, *D* is a root node.

The experimenter decides which treatment a sample receives rather than, e.g., the state of the cells, so nothing unmeasured influences assignment. Randomization would be sufficient, but it is not necessary. External assignment is enough.

Under Assumption 1, the parameters (*α*^6^*, β*^6^) in model (1) describe the total causal effect of treatment on protein expression at 6 hours. Writing do(*D* = *d*) for an intervention that administers the concentrations *d*, see [5], these parameters describe how

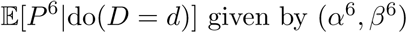

changes with *d*, where *P^t^* is the vector of differential protein expressions at time *t* relative to the untreated baseline of the same cell line. The group p-values for (*α*^6^*, β*^6^) therefore test whether the intervention changes expression of protein *r* at 6 hours.

The same argument applies at 24 and 48 hours if the later expression is the response of model (1). Under Assumption 1 alone, the parameters (*ᾱ^t^, β̄^t^*) describe how E[*P^t^* | do(*D* = *d*)] changes with *d* for *t* ∈ {24, 48}, which is the total causal effect at time *t*.

Working with differential expression adjusts for cell-line effects. The baseline is measured before treatment and depends only on the cell line, so it is not a consequence of *D*, and subtracting it cannot open a path between treatment and expression. Two things follow from this step. We assume cell-line effects are additive on the log scale, and we average the reported effects over the 18 cell lines, since we do not model drug-by-cell-line interaction. A drug acting strongly in a few lines and weakly in the rest therefore shows a diluted effect.

The models at 24 and 48 hours condition on protein expression at the preceding time point, which the treatment itself affects. Their *α* and *β* parameters are direct effects, meaning the part of the treatment effect that does not pass through the modeled earlier expression. They describe how

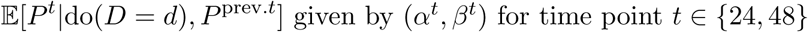

changes with *d*, for *t* ∈ {24, 48}. Because conditioning on the earlier expression could in principle open up confounding paths, reading them causally requires a second assumption.

#### Assumption 2

Protein expressions at two consecutive time points (6 and 24 hours, or 24 and 48 hours) have no unmeasured common causes.

Assumption 2 is considerably stronger than Assumption 1 and may fail. Proteins outside the measured set, metabolites, and other molecular processes can influence expression at more than one time point, which is what *U* represents in Figure 5. Where Assumption 2 fails, conditioning on the earlier expression can produce an apparent association between treatment and later expression even when no direct effect exists, because the earlier expression is a common consequence of treatment and of *U*. We therefore treat the total effects at all three time points as the best-supported causal claims, and the direct effects at 24 and 48 hours with less certainty.

The *γ* parameters relate expression of one protein at an earlier time point to expression of another at the next. Their direction follows the temporal ordering of the measurements and needs Assumption 2 to be causal, since *U* influences both endpoints of such an edge. We do not interpret these edges mechanistically. They help generate hypotheses about which proteins move together over time after perturbation and suggest which candidates to follow up experimentally.

In biological terms, the parameters of models (1) and (2) read as follows. The single-drug effects *α* describe the change in a protein’s abundance caused by administering a drug, compared with not administering it under otherwise identical conditions. The interaction effects *β* describe how far the change under a drug pair departs from the sum of the changes under its two drugs. Because the experimenter controls the treatment, these are effects of an intervention and not associations between treatment and abundance, under Assumption 1 for the total effects and additionally under Assumption 2 for the direct effects. The temporal dependencies *γ* make a weaker claim. A significant *γ* says that differential expression of one protein is followed by differential expression of another, after removing the residual drug effects, which is consistent with a regulatory relationship but also with both responding to something unmeasured.

### Nonlinear effects and linear approximation

The linear models described above in equations (1)-(2) can be interpreted as best linear approximations to potentially nonlinear true relationships. While such approximations may not capture all nonlinear effects, the resulting p-values remain valid, under some assumptions, for testing the null hypothesis of no direct causal effect. Specifically, these assumptions concern the distribution of covariates: for example, for multivariate Gaussian distributions, if there is no relation (linear or nonlinear), the linear approximation will not produce false associations or causal statements. Conversely, the method is conservative, as certain nonlinear relationships may remain undetected. For details, see [41]. Thus, the linear model equations in (1)-(2) typically yield correct causal inference statements: a significant effect is also significant in a nonlinear model, but we may not detect some nonlinear effects. Making the model equations in (1)-(2) more nonlinear leads to unfavorable ratios of parameters (to be estimated) and sample size. While equation (1) has up to 244 parameters, the equations in (2) have up to 5’322 parameters, and modeling nonlinearities in such a high-dimensional setting is challenging and much more unstable.

### 4.1 Statistical validation for the IC50 phenotype

We consider here the modeling and estimation of the response of IC50 values. We model the IC50 response as a potentially nonlinear function of protein expression levels, ideally invoking only the causal (but unknown) proteins. For this task, we proceed as follows.

#### Random Forests with invariance regularization

The IC50 values come from two sources. For the single drugs, Sun et al. [28] measured them in a separate cell viability assay, described in the Methods of that study. For the drug combinations, we use the combination IC50 values of [42] for the cell lines and drug pairs studied here. Both enter the analysis as log2 values in µM, and for the 18 cell lines (16 triple-negative and 2 non-triple-negative breast cancer) we obtain *N* = 1599 values, which we stack in a response vector ∈ ℝ^1599^. The two sources differ in laboratory and protocol, so IC50 values are comparable within single drugs and within combinations, while differences between the two groups partly reflect the source. We model *IC*50*_i_* with the aggregated differential log-protein expression 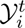 at all three time points for the corresponding experimental condition *C_i_* and *D_i_*, as described in the subsection on “Structure of the dataset and notation”. Note that the samples, i.e., each sample unit *i*, are unpaired because they come from different experiments measuring IC50 or protein expression. This results in the model

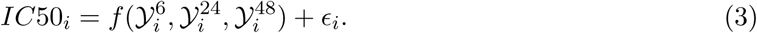

We estimate the potentially nonlinear function *f* (*., ., .*) by Random Forests [43] but with an added causality-inspired invariance regularization, see (4) below. By treating different drug perturbations as different experimental settings, invariance regularization aims to make residual distributions similar across environments. This leads to provable robustness guarantees in so-called anchor regression settings [44], and, under idealized assumptions, such as those from instrumental variables regression [45–47], it infers causal relations. Invariance regularization can be used with almost any machine learning algorithm [48, 49] by minimizing the anchor objective

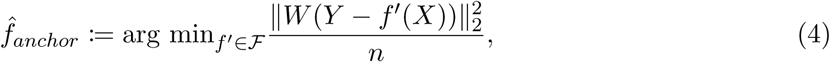

where 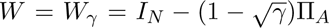 with Π*_A_* being the linear projection operator onto *A* and *γ* controlling the strength of invariance regularization.

In our case, *A* = *D* (the values for the different drug concentrations which are used for treatment), *Y* = *IC*50 (the vector of the observed IC50 values), and 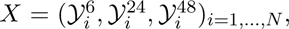 where we concatenate the aggregated differential log-protein expression column-wise. We choose Random Forests for their power relative to their simplicity and their lower sample-size requirements compared to more complex models, such as deep learning. Ulmer et al. [50] minimize a transformed version of the residuals in the context of spectral deconfounding. We use their software to minimize Equation (4), resulting in Anchor Forests, see also [51].

The regularization strength *γ* controls the trade-off between fitting *IC*50 well and restricting the used signal and the used proteins to that part that stays similar across different treatments. Here, we want to choose *γ* to predict *IC*50 as well as possible, even for a new drug treatment consisting of a drug combination.

The 59 drug combinations fall into eight groups, each formed by the combinations that share one of eight drugs, which is combined with a range of other drugs. The eight drugs are Olaparib, Lapatinib Ditosylate Hydrate, Vorinostat, Temozolomide, Docetaxel, Gemcitabine Hydrochloride, Cisplatin, and 5-Fluorouracil, and a combination of two of them belongs to the group of the drug listed first in its experimental annotation. We now approximate the shift to a new unseen drug combination using these natural environments. So instead of optimizing predictive power for a new drug combination, we optimize it for shifts as strong as those observed across these eight environments. In particular, we want to be worst-case optimal and perform as well as possible, even if a new double-perturbation shifts as much as the most extreme environment.

To choose the right amount of invariance regularization *γ* for this goal, we use these environments for an out-of-environment cross-validation scheme.

We repeatedly estimate Anchor Forests for training with all single-drug perturbations and seven of the eight environments, using the drug concentrations as Anchor variables and different *γ* values, and evaluate the estimated models on the left-out environment. Figure 6 shows the out-of-environment mean squared error for *IC*50 for the eight environments and varying *γ*. The thick black line corresponds to *γ* = 1, resulting in standard Random Forests. Points show the error for each *γ*, and lines are LOESS-smoothed [52] for display.

**Figure 6:**
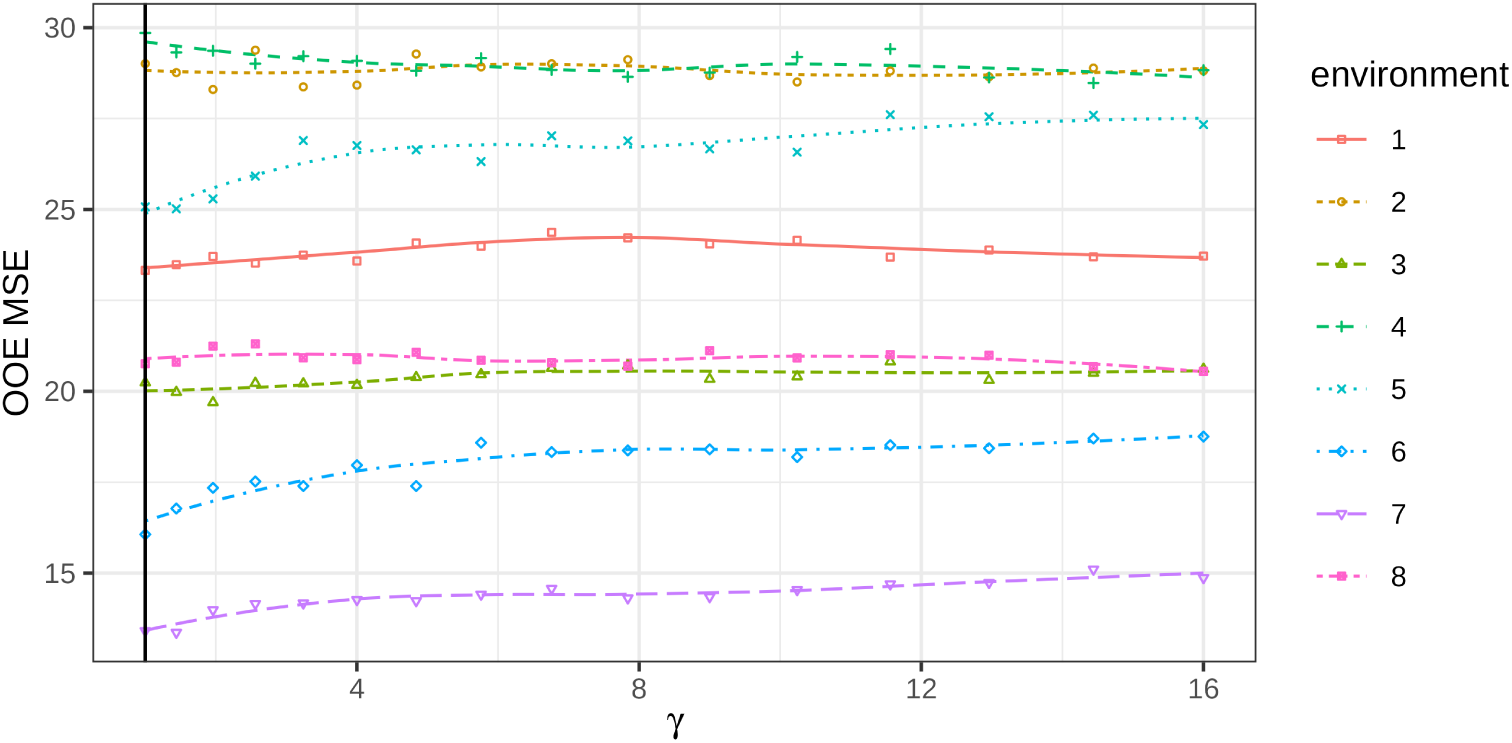
The out-of-environment error changes little with the invariance regularization for *γ* ≥ 1. Shown are the out-of-environment mean squared errors (OOE MSE) for the eight left-out environments, where the Anchor Forest is fitted on all data except the environment it predicts. Points are the errors for each *γ*, and lines are LOESS-smoothed for display. The black vertical line at *γ* = 1 marks the standard Random Forest without invariance regularization.

We restrict *γ* to values of at least 1, the range in which anchor regression provides robustness against shifts in the span of the anchor variables [44]. As the most difficult environment, we take the one with the largest out-of-environment mean squared error of the standard Random Forest, which is the group of combinations with Temozolomide (environment 4). Its out-of-environment error is smallest at *γ* = 14.44, an improvement of 4.6% over the standard Random Forest, and we use this value for the remainder of the analysis. Because *γ* is chosen on this environment, the gain is an optimistic estimate. Over *γ* ≥ 1, the error of this environment varies only between 28.48 and 29.85, and the other environments show similarly flat curves, so a broad range of *γ* performs comparably. The standard Random Forest improves the out-of-environment error over a constant prediction by the training mean by 2.5% to 21.8% across environments. The proteome therefore carries moderate predictive signal for the IC50 of new drug combinations, and the invariance regularization adds a small gain in the most difficult environment.

In a second step, we re-estimate an Anchor Forest on all data (single drug perturbations and all eight environments) with *γ* = 14.44. We then analyze the estimated invariant model and the key proteins it identifies.

Eight proteins have variable importance above 2 in the invariant model, separating them from the remaining predictors in the regularization path (Figure 8). Six of them are measured at 48 hours (TPX2, IFI30, THBS1, RACK1, SNRPD2, and C3), CKS1B at 24 hours, and ITGA3 at 6 hours. The two most important are TPX2 and IFI30 at 48 hours. Partial dependence analysis of the estimated Anchor Forest (Figure 7) shows a step for each protein. Treated conditions in which TPX2 rises at 48 hours, mostly from below the detection limit at baseline, have a higher predicted IC50, and treated conditions in which IFI30 decreases, in part to below the detection limit, have a lower predicted IC50. Changes beyond about 2 in absolute value arise when a protein crosses the detection limit between baseline and 48 hours, so their size reflects the imputed floor rather than the abundance. Both steps are small, about 0.05 on the log2 scale, corresponding to about a 4% change in IC50.

**Figure 7:**
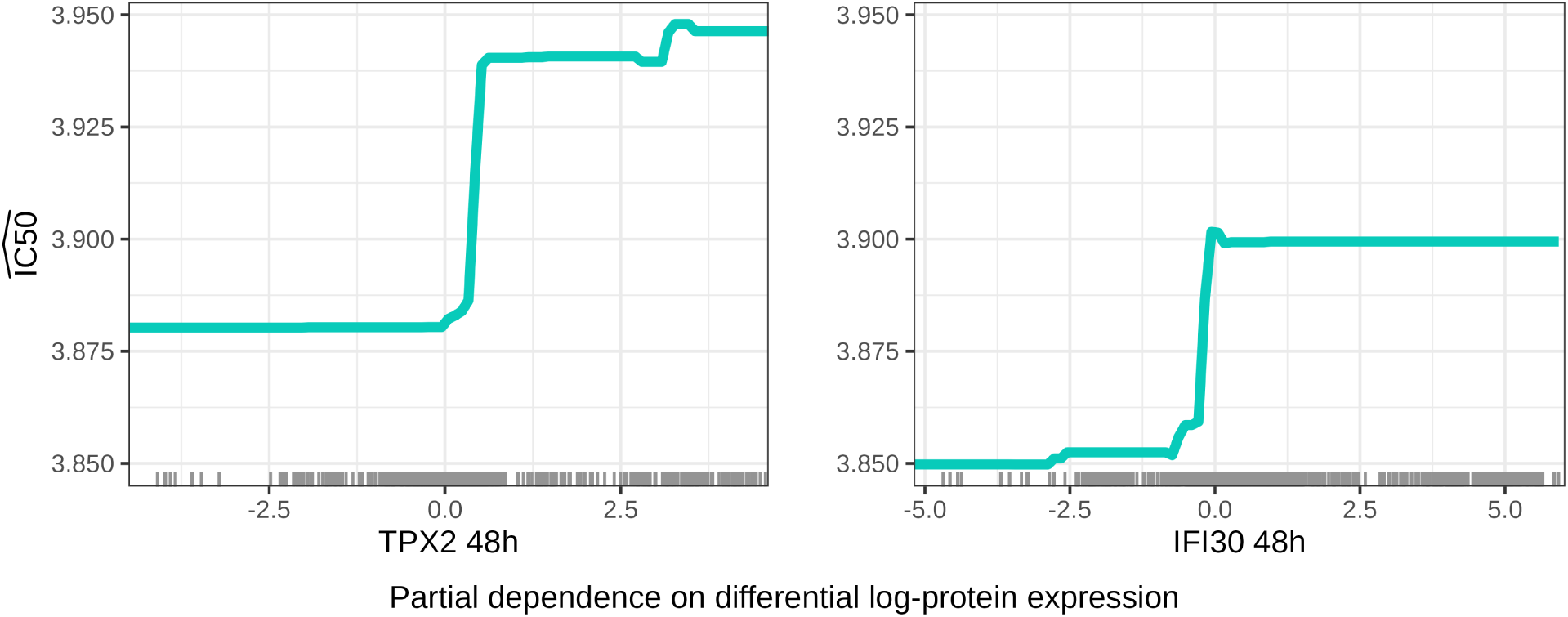
TPX2 becoming detectable at 48 hours raises the predicted IC50 and a drop of IFI30 lowers it, both by about 4%. Shown are partial dependence plots for TPX2 and IFI30 at 48 hours, the two most important proteins in the Anchor Forest refitted with *γ* = 14.44. For all observations, the predictor of interest is varied, yielding *N* predictive functions, and the bold line averages over the *N* functions.

**Figure 8:**
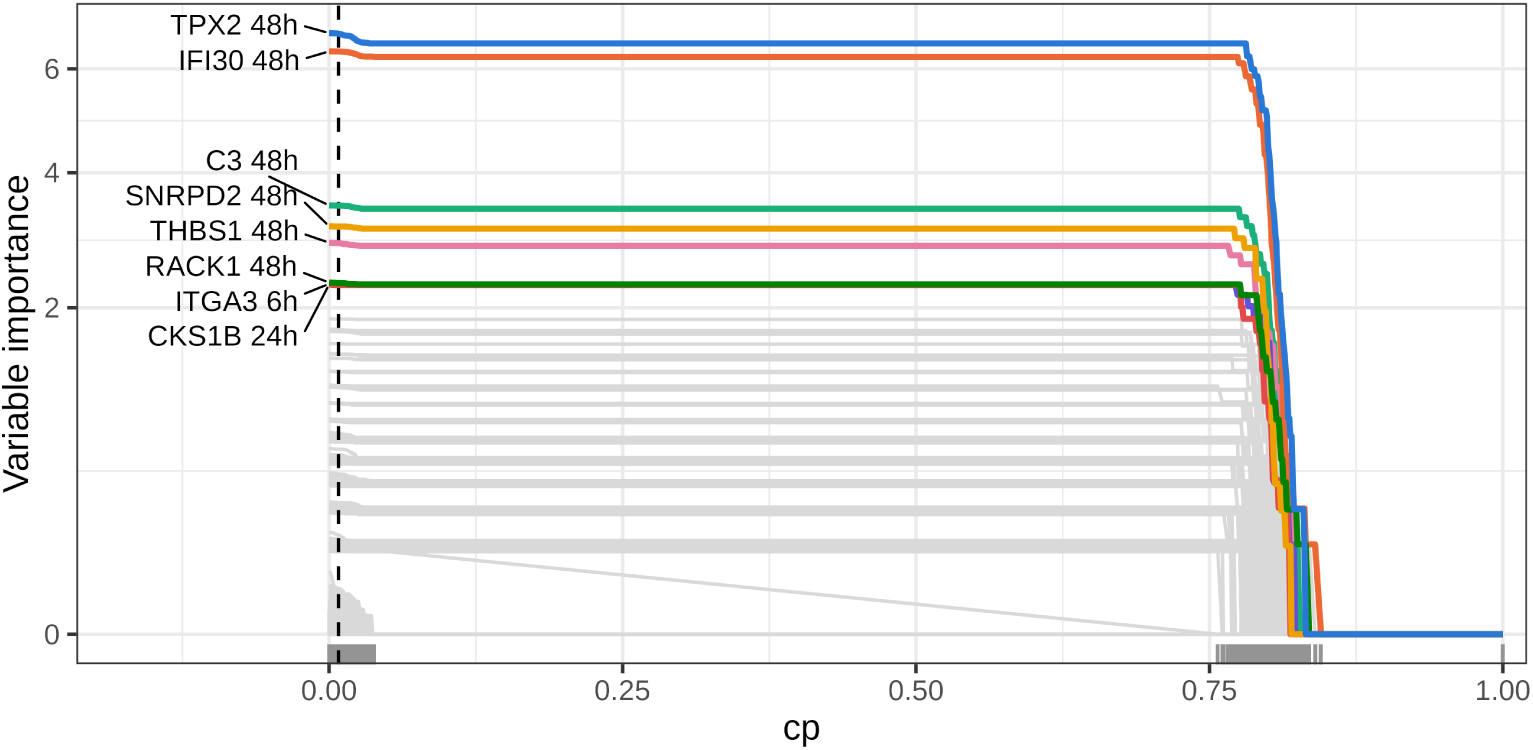
Regularization path of refitted Anchor Forest with *γ* = 14.44. The eight selected proteins are drawn in color and labeled with their measurement time point, and all other predictors are drawn in gray. The figure shows the variable importance of different predictors (proteins at different time points) as pruning increases (cost complexity *cp*) in the Forest (*cp* = 0 means no pruning), with variable importance on a square-root scale. Variable importance is the decrease in loss from splits using a specific predictor. The vertical line shows the *cp* value with minimal out-of-bag error, which is close to 0 (no pruning).

Two of the eight proteins belong to the proliferation program of the cell cycle. TPX2 activates Aurora A and nucleates microtubules during spindle assembly, and high nuclear TPX2 is associated with high-grade, estrogen- and progesterone-receptor-negative status, and poor outcomes in breast cancer [53]. CKS1B encodes Cks1, which interacts with cyclin-dependent kinases and is an essential cofactor for the SCF^Skp2^-mediated degradation of the inhibitor p27, and its overexpression is associated with loss of differentiation in breast tumors [54]. TPX2, which becomes detectable 48 hours after treatment, is associated with a higher predicted IC50, consistent with cells that maintain or resume proliferation under treatment being less sensitive to cytotoxic drugs, most of which act on dividing cells.

Two proteins belong to the translation and splicing machinery. RACK1 is a scaffolding protein that binds the 40S ribosomal subunit and signaling kinases such as PKC and Src [55], and high RACK1 expression predicts poor outcomes in breast cancer [56]. SNRPD2 is one of the Sm proteins that form the core of the spliceosomal snRNPs [57]. Both can be read as markers of the overall biosynthetic state of a cell under treatment.

The remaining four proteins act at the cell surface or in the extracellular space. IFI30 encodes the lysosomal thiol reductase GILT, an interferon-*γ*-inducible enzyme of antigen processing. High IFI30 expression is associated with poor prognosis in breast cancer, and its knockdown reduces the proliferation of breast cancer cells [58]. Lower IFI30 after treatment, including its loss below the detection limit, is associated with lower predicted IC50. THBS1 encodes thrombospondin-1, a glycoprotein that inhibits angiogenesis [59]. C3 is the central component of the complement system, which tumor cells produce locally and which modulates tumor growth and the immune microenvironment [60]. ITGA3 encodes the integrin *α*3 subunit, and integrins link cells to the extracellular matrix, including the laminins of the basement membrane, and transmit signals across the plasma membrane in both directions [61]. These proteins describe a cell’s interaction with its environment rather than its division.

Together, the eight proteins cover proliferation, biosynthesis, and the interaction with the microenvironment, three aspects of the cellular state that can influence drug sensitivity. In the temporal protein network (Figure 3(a)), RRM2 at 24 hours is an antecedent of TPX2 and CKS1B at 48 hours, and the biological plausibility of this link is discussed in S1 Appendix, Section B.

Figure 8 shows the regularization path of the estimated Anchor Forest. It shows variable importance (i.e., how much a protein reduces loss) as the forest is pruned. This provides a tool to analyze the path from a very simple random forest with very simple trees (stumps only at *cp* = 1) to a very complex random forest using a large number of proteins.

In Figure 8, the eight proteins above stand out from the remaining predictors by a visible gap in variable importance with stronger pruning. We use these eight proteins for downstream exploration with Drug-Prot (Section 2).

From a translational perspective, these findings suggest a potential framework for leveraging early proteomic responses to drug perturbations. For example, in an ex vivo or experimental precision-medicine setting, baseline protein expression could be assessed before treatment, followed by measurement of these predictive proteins at post-treatment time points. The resulting profiles could then be used to estimate drug sensitivity using the proposed Anchor Forest model.

TPX2 and IFI30 may be candidates for evaluating a reduced marker panel, but the predictive performance of such a panel remains to be established. Monitoring changes in the abundance of these two proteins 48 hours after drug exposure, one marker of proliferation and one of the interferon-inducible lysosomal response, could provide a coarse readout of cellular drug responsiveness. While such applications remain speculative and would require substantial validation, they highlight how proteomic dynamics after treatment may inform treatment stratification strategies.

### 4.2 Use of AI tools

We used Claude (Anthropic) to edit and draft parts of the manuscript, support the literature review, and assist with writing and reviewing analysis, figure, and application code. The schematic drawings in Figures 1(a), 1(b) and 5 were drawn by R scripts written with Claude. All statistical modeling and corresponding choices were made by the authors, and all code was reviewed and run by the authors. We take full responsibility for the content of this article.

## 5 The Drug-Prot platform

Drug-Prot is implemented as an R Shiny dashboard and is publicly hosted at https://markus-ulmer.ch/drugprot/. The source code is available at https://github.com/markusul/DrugProt. All statistical evidence (approximately 55 million p-values across 5’078 proteins, 122 treatments, and three post-treatment time points) is precomputed and bundled with the application, so the user never needs to access or download the underlying proteomic dataset.

### Query inputs

Users specify a set of proteins of interest in one of three ways: (i) typing or selecting from a searchable picker of all 5’078 measured proteins, (ii) clicking one of two example buttons, which load the two most IC50-predictive proteins (TPX2 and IFI30) or all eight IC50-predictive proteins of Section 2, or (iii) uploading a plain-text file with one protein identifier per line. Proteins are referred to throughout by their HGNC gene symbol, with co-quantified protein groups joined by a slash (e.g. CALM1/CALM2/CALM3).

Two additional settings control the inference: the family-wise or false-discovery significance level *α* (default 0.05), and separate multiple-testing corrections for drug effects (Holm by default) and for protein-to-protein temporal dependencies (Benjamini–Hochberg by default). The tests are always two-sided for no effect versus non-zero effect. All standard correction methods from `R` [62] are available. The set of analyzed time points can be restricted to any subset of {6, 24, 48} hours.

### Drug-effect outputs

For the chosen protein set, Drug-Prot returns the minimum corrected p-value over the set and the chosen time points for each of the 63 single drugs and each of the 59 experimentally available drug pairs. These p-values are shown as an interactive 63 × 63 heatmap where the diagonal encodes single-drug evidence and the off-diagonal encodes pairwise interaction evidence. For 24 and 48 hours, users choose whether the p-values refer to the direct effects in model (2) or to the total effects in model (1), which coincide at 6 hours. Unmeasured drug pairs are shown in cyan, outside the p-value color scale. Two companion tables list, in the order of increasing p-values, the significant single drugs and the significant drug pairs. We additionally report the average treatment effects and their significance on all proteins in the set.

### Network outputs

For the queried protein set, Drug-Prot returns the directed temporal dependency subgraph, based on significant findings, linking the queried proteins to other proteins in the dataset. Two visualizations are provided. A *temporal graph* in which each node is a (protein, time point) pair and edges go from *t* = 6 to *t* = 24 and from *t* = 24 to *t* = 48 whenever the corresponding *γ* parameter in model (2) is significant at the user-chosen level, and a *summary graph* that collapses the temporal graph across time. Both graphs are rendered as zoomable, draggable, force-directed networks with arrows. In the *temporal graph*, the edges are colored by positive or negative effects. A companion table reports, for each queried protein, the numbers of parents and children outside the query set, along with their identities.

### Downloads

Users can download (i) the corrected drug-effect p-values for the queried protein set and the chosen effect type as a CSV, including the estimated effect and its sign when a single protein is queried, (ii) the corrected p-values and signs of every protein–protein edge involving the queried set, by time-point transition, as a CSV, and (iii) the summary and temporal graphs as standalone, interactive HTML files that can be embedded in supplementary material.

### Scope of analyses

Because all p-values are precomputed, the query response is near-instantaneous regardless of the size of the protein set. The multiple-testing correction is applied only to the queried set and the selected time points, and thus, analyses focused on a small, biologically motivated set of proteins yield substantially greater power than proteome-wide queries.

Drug-Prot is currently a point-and-click web application. Programmatic or batch access is not yet available via a dedicated API, but the complete precomputed evidence used by the application can be downloaded from the platform for users who wish to run large-scale or scripted queries offline.

## Data/Code availability

The perturbation proteomics dataset is published in [28]. The processed protein matrix and sample annotation are available at https://db.prottalks.com, and the raw mass spectrometry data is deposited in ProteomeXchange via iProX under accession IPX0007409000. The sample selection, protein identifiers, and single-drug IC50 values used in this study, together with the full statistical evidence underlying the application, are deposited at Zenodo (DOI: 10.5281/zenodo.21508012). The combination IC50 values are taken from the original anchored screen of [42], available on Figshare (DOI: 10.6084/m9.figshare.16843597). Source code for the query application is available at https://github.com/markusul/DrugProt and archived at Zenodo (DOI: 10.5281/zen-odo.21510303). Analysis code, which reproduces all results from the public data, is available at https://github.com/markusul/DrugProt-Paper and archived at Zenodo (DOI: 10.5281/zen-odo.21509421).

## Acknowledgements

We thank every member of the scientific community with whom we had fruitful discussions about Drug-Prot. We especially thank Michael Vollenweider for his helpful questions, comments, and inputs.

## Supporting information

### S1 Appendix. Supporting information

Contains (A) description of the perturbation proteomics dataset, (B) biological plausibility of the temporal dependencies among the selected proteins, and (C) a case study on drugs that change the abundance of their target.

