## Supplementary material for "Drug-Prot: A query system for statistical inference of drug effects and interactions in dynamic proteomic networks": S1 Appendix

#### A The perturbation proteomics dataset

All statistical evidence presented in `Drug-Prot` is derived from the large-scale perturbation proteomics dataset of [1], to which we refer for the full experimental, mass spectrometry, and data-processing details. Here we summarize the dataset features relevant to the models in (1) and (2) and establish the correspondence to our notation in Section 4.

**Design.** The dataset profiles the proteomic response of breast cancer cell lines to clinically relevant drug perturbations, resolved over time. It is organized along four axes: cell line, perturbation (single drug or drug pair), post-treatment time point, and biological replicate. Concretely, Sun et al. [1] treated 18 breast cancer cell lines (sixteen triple-negative and two non-triple-negative) with a panel of 63 FDA-approved small-molecule drugs spanning chemotherapeutic, targeted, and hormonal classes, and additionally administered selected two-drug combinations. Protein expression was measured at 6, 24, and 48 hours after treatment, in three biological replicates per condition, with untreated controls providing a pre-treatment baseline. The treated cell lines consist of BT20, BT549, DU4475, HCC1143, HCC1187, HCC1395, HCC1806, HCC1937, HCC38, HCC70, HS578T, MCF7, MDA-MB-231, MDA-MB-436, MDA-MB-453, MDA-MB-453(ATCC), MDA-MB-468, and T47D. The drugs used were Cisplatin, Docetaxel, Lapatinib Ditosylate Hydrate, Gemcitabine Hydrochloride, Olaparib, Temozolomide, Vorinostat, 5-Fluorouracil, Afatinib Dimaleate, Paclitaxel, Sorafenib Tosylate, Alpelisib, Indocyanine Green, Ixabepilone, Methotrexate, Epirubicin Hydrochloride, Sonidegib Diphosphate, Doxorubicin Hydrochloride, Tucatinib, Cobimetinib Fumarate, Bosutinib Monohydrate, Entrectinib, Talazoparib, Abemaciclib, Neratinib Maleate, Letrozole, Niraparib Tosylate, Enzalutamide, Pazopanib Hydrochloride, Phenylephrine Hydrochloride, Toremifene citrate, Melphalan, Hydrocortisone, Ribociclib Succinate, Vandetanib, Decitabine, Sunitinib Malate, Erlotinib Hydrochloride, Azacytidine, Fulvestrant, Bexarotene, Verteporfin, Aminolevulinic Acid Hydrochloride, Anastrozole, Estradiol,

Oxaliplatin, Pemetrexed Disodium Hydrate, Everolimus, Zoledronic Acid Hydrate, Raloxifene Hydrochloride, Vincristine Sulfate, Capecitabine, Irinotecan Hydrochloride, Carboplatin, Cyclophosphamide, Medroxyprogesterone Acetate, Chlorambucil, Selumetinib sulfate, Dasatinib Hydrate, Eribulin Mesylate, Trilaciclib Dihydrochloride, N-Descyclopropanecarbaldehyde Olaparib, and Lurbinectedin. We index the 63 drugs by $J = \{1, \ldots, 63\}$ and the cell lines by $\{1, \ldots, 18\}$, consistent with the drug-concentration matrix $D$ and the cell-line index $C_i$ introduced in Section 4. The individual drugs are cataloged in [1].

**Perturbation conventions.** Single-drug experiments were administered at a fixed concentration of 10 µM, so that exactly one entry of the corresponding row of $D$ is non-zero. Combination experiments pair two drugs at concentrations between 0.00025 and 30 µM, so that exactly two entries are non-zero. Our analysis covers the 63 single drugs together with the 59 experimentally measured drug pairs, for a total of 122 treatments referred to throughout.

**Scale and acquisition.** The acquisition campaign of [1] produced about $1.6 \times 10^4$ DIA mass-spectrometry runs and more than $3.8 \times 10^7$ individual protein measurements, quantifying several thousand protein groups per condition. To our knowledge, this makes it the largest perturbation proteomics resource of its kind to date. After the quality control of [1] and the filtering described below, the data entering our analysis comprise 5'078 proteins, measured across 4'625 samples at $t = 6$, 4'649 at $t = 24$, and 4'540 at $t = 48$ hours. These per-time-point counts are the perturbed samples in models (1) and (2), and at 24 and 48 hours they include only samples whose experimental condition was also measured at the preceding time point. The larger total run count also includes baseline ($t = 0$), control, and quality-control acquisitions.

**Sample selection and missing values.** The analysis uses the protein intensity matrix and the sample annotation of the public release of [1], together with the list of selected samples deposited on Zenodo (DOI: 10.5281/zenodo.21508012). Of the 1'117 untreated controls in the release, we selected the 840 that meet the criteria of complete biological triplicates and a full factorial design. The intensity matrix also contains 158 technical replicates that are not part of the sample annotation, 11 of these are among these controls, and we exclude them. The baseline for each cell line is the median across its untreated controls and the double-baseline samples at 0 hours, for 931 samples in total. The double-baseline samples at 6, 24, and 48 hours receive no drug and enter models (1) and (2) with $D_i = 0$ as the time-matched reference.

Missing protein intensities are treated as values below the detection limit and set to 80% of the smallest observed intensity of the protein before the log transformation. We keep proteins with more than ten distinct values, which leaves 5'392 proteins. We further keep proteins whose differential expression differs from its most frequent value in at least ten samples at every time point, in the samples and in the lagged design, which leaves 5'078 proteins. For models (1) and (2), the values at this detection floor are replaced within each cell line by the conditional mean of a normal distribution fitted to the observed values of that protein and cell line and truncated at the smallest observed value. This imputation is deterministic, so all results are reproducible from the public data.

**From measurements to model inputs.** As protein quantification is destructive, each sample is observed at a single time point. The data is therefore *unpaired* across $t \in \{6, 24, 48\}$. Whenever a protein expression at one time point is modeled in terms of expressions at an earlier time point (as in (2)), the earlier values must therefore be replaced by the within-condition aggregate $\mathcal{P}^t_{r:i}$ defined in Section 4, the median log-expression over all samples sharing the experimental condition $(C_i, D_i)$. Second, all expression levels enter as *differential* quantities $\mathcal{Y}^t_{r:i}$ relative to the matched untreated baseline of the same cell line, which implicitly adjusts for cell-line effects. The drug structure used in [1] (molecular fingerprints and physicochemical descriptors) is not used by `Drug-Prot`, which depends on the dataset only through the perturbation matrix $D$, the cell-line index, and the differential protein expressions.

### B Biological plausibility of the temporal dependencies among the selected proteins

The following interpretations aim to assess biological plausibility rather than to establish direct molecular mechanisms. The edges are statistical dependencies in the sense of model (2), and their direction follows the temporal order of the measurements.

- **RRM2 as an antecedent of TPX2 and CKS1B**

  RRM2 measured at 24 hours is an antecedent of TPX2 and of CKS1B measured at 48 hours, with positive coefficients for both edges. RRM2 is the regulatory subunit of ribonucleotide reductase, the enzyme that produces the deoxyribonucleotides for DNA synthesis [2], and its expression rises at the onset of S phase [3]. TPX2 acts in spindle assembly during mitosis, and CKS1B interacts with cyclin-dependent kinases and promotes the degradation of p27 [4, 5]. Cell-cycle transcript profiling has associated RRM2 with S phase and TPX2 and CKS1B with G2/M [6]. The positive temporal associations observed here are therefore compatible with a shared proliferation-related response to treatment. However, the measurements were obtained from separate cell populations at each time point and do not establish progression from S phase at 24 hours to G2/M at 48 hours. These associations may reflect differences in proliferative state or cell-cycle composition across conditions, rather than direct regulation of TPX2 or CKS1B by RRM2.

  This dependence connects to the drug results of the use case. Gemcitabine inhibits ribonucleotide reductase and is incorporated into DNA [7], and increased RRM2 expression is associated with Gemcitabine resistance in cancer cells [8]. The temporal associations of RRM2 with TPX2 and CKS1B suggest a testable hypothesis that nucleotide metabolism and proliferative state may contribute to the protein responses observed under Gemcitabine treatment and its combinations. These associations do not establish that RRM2 mediates the drug effects or interactions.

- **RACK1 and its downstream proteins**

  RACK1 is an antecedent of 20 proteins, all with positive coefficients, among them PCNA and the chaperonin subunit CCT5 from 6 to 24 hours, and the deacetylase HDAC1 and the proteasome subunits PSMD14, PSMD4, and PSMB2 from 24 to 48 hours. RACK1 is a scaffolding protein that binds the 40S ribosomal subunit and signaling kinases [9]. Its downstream proteins span replication, protein folding, and protein degradation, which together describe a cell's biosynthetic activity. We read these edges as a common response of the biosynthetic program to treatment, in which the level of RACK1 indicates whether a cell has maintained its capacity for protein synthesis, rather than as individual regulatory relations.

- **Self-dependencies**

  THBS1, RACK1, C3, TPX2, and IFI30 depend on their own expression at the preceding time point, with the strongest dependence for THBS1 ($p = 9 \times 10^{-18}$ from 24 to 48 hours). These edges reflect the persistence of treatment-induced expression changes over time and serve as a positive control for the temporal model, since a protein whose change at 24 hours does not predict its change at 48 hours would indicate either a transient response or a lack of signal.

- **Summary**

  The temporal dependencies among the selected proteins organize around two hubs. Earlier RRM2 abundance is associated with subsequent levels of the proliferation-related proteins TPX2 and CKS1B, and earlier RACK1 abundance with subsequent levels of proteins of replication, folding, and degradation. Both hubs are consistent with a reading of the eight proteins as markers of whether a cell continues its proliferative and biosynthetic program under treatment. These links remain hypotheses generated by the statistical analysis and require experimental validation.

### C Case study on drugs that change the abundance of their target

Some drugs change the amount of the protein they act on, and pharmacology predicts the direction. An inhibitor blocks the activity of its target, not its production, and for TYMS and DHFR the amount rises under inhibition, because each enzyme limits its own amount through the repression of its mRNA translation and through its turnover, and the inhibitor-bound enzyme escapes both. Methotrexate binds dihydrofolate reductase (DHFR) and prevents the enzyme from binding its own mRNA, relieving repression of its translation so DHFR accumulates [10], an increase first observed in cultured cells by [11]. 5-Fluorouracil raises thymidylate synthase (TYMS), whose protein represses the translation of its own mRNA until the 5-Fluorouracil metabolite binds [12] and whose inhibitor-bound form is protected from degradation in cells [13]. Pemetrexed inhibits both TYMS and DHFR [14], so we expect both enzymes to increase after it, as TYMS does after 5-Fluorouracil and DHFR after Methotrexate. Decitabine is incorporated into DNA and traps DNA methyltransferase 1 (DNMT1), and the adducts trigger the proteasomal degradation of the free enzyme [15], whereas Azacytidine is a ribonucleoside analog that reaches DNA only after further conversion [16], so we expect a weaker effect. Camptothecins, the class of the active metabolite of Irinotecan, trap topoisomerase I (TOP1) on DNA and trigger its ubiquitin-dependent degradation [17], and topoisomerase II poisons trap topoisomerase II$\beta$ (TOP2B) in covalent complexes that the proteasome processes, shown for teniposide [18] and proposed for doxorubicin [19]. Lurbinectedin stalls elongating RNA polymerase II and triggers the degradation of its phosphorylated form (POLR2A) [20]. Glucocorticoids induce FKBP5 through the glucocorticoid receptor [21], and Medroxyprogesterone Acetate binds and signals through this receptor, although with little transactivation in lymphocytes [22]. Drugs that bind their target without trapping it should leave its amount unchanged. We expect this for the kinase inhibitors of EGFR, and for the PARP inhibitors, whose trapping of PARP1 on DNA is rapidly reversible [23]. Capecitabine is activated in three steps, the first of which occurs almost exclusively in the liver, and it is only weakly cytotoxic in cultured cells [24], so no change in TYMS is expected either.

We queried the nine targets DHFR, TYMS, DNMT1, TOP1, TOP2B, POLR2A, FKBP5, PARP1, and EGFR in `Drug-Prot` with Holm correction at $\alpha = 0.05$ across the set and all three time points, with the direct drug effects at 24 and 48 hours. At the level of the set, 57 of the 63 single drugs are significant, 56 of them through POLR2A, which changes 6 hours after drug perturbation, upward for all but the two taxanes, as part of the shared early response described in the main text. We proceed by reading the results per target and time point separately. Table 1 lists them for the 21 drug-target pairs with a prediction, 12 with a predicted change and 9 predicted to stay unchanged.

Table 1: Drug-Prot recovers five of the twelve predicted changes in target abundance with the right direction and time course, and none of the nine drug-target pairs predicted to stay unchanged shows a significant change at 24 or 48 hours. Each row is a drug-target pair, and the EGFR row combines five pairs. Holm-corrected p-values over the nine queried proteins, 122 treatments, and three time points, with the direction of the estimated effect where the p-value is below 0.05. At 24 and 48 hours, the p-values refer to the direct effects of model (2). The five EGFR inhibitors are Afatinib, Erlotinib, Lapatinib, Neratinib, and Tucatinib.

| Target | Drug | Expected | 6 h | 24 h | 48 h |
|---|---|---|---|---|---|
| DHFR | Methotrexate | up | 0.12 | $\uparrow 5.1 \times 10^{-4}$ | $\uparrow 2.3 \times 10^{-8}$ |
| DHFR | Pemetrexed | up | 0.19 | 1 | $\uparrow 1.9 \times 10^{-7}$ |
| TYMS | 5-Fluorouracil | up | 1 | 1 | $\uparrow 1.7 \times 10^{-3}$ |
| TYMS | Pemetrexed | up | 1 | 0.17 | 1 |
| DNMT1 | Decitabine | down | 1 | 0.12 | $\downarrow 0.048$ |
| DNMT1 | Azacytidine | down | 1 | 1 | 1 |
| TOP1 | Irinotecan | down | 1 | $\downarrow 1.4 \times 10^{-7}$ | $\downarrow 2.0 \times 10^{-6}$ |
| TOP2B | Doxorubicin | down | 1 | 1 | 1 |
| TOP2B | Epirubicin | down | 1 | 1 | 1 |
| POLR2A | Lurbinectedin | down | 0.32 | 1 | 1 |
| FKBP5 | Hydrocortisone | up | 1 | 1 | 1 |
| FKBP5 | Medroxyprogesterone | up | 1 | 1 | 0.67 |
| TYMS | Capecitabine | none | 1 | 1 | 1 |
| PARP1 | Olaparib | none | 1 | 1 | 1 |
| PARP1 | Niraparib | none | $\uparrow 1.3 \times 10^{-5}$ | 1 | 1 |
| PARP1 | Talazoparib | none | $\uparrow 0.03$ | 1 | 1 |
| EGFR | five EGFR inhibitors | none | 1 | 1 | 1 |

`Drug-Prot` recovers five of the twelve predicted changes. DHFR increases after Methotrexate at 24 and 48 hours and after Pemetrexed at 48 hours, TYMS increases after 5-Fluorouracil at 48 hours, DNMT1 decreases after Decitabine at 48 hours, and TOP1 decreases after Irinotecan at 24 and 48 hours. At 24 and 48 hours, no other single drug has a significant effect on these four targets, so the significant effects on DHFR come only from its two inhibitors Methotrexate and Pemetrexed, on TYMS from 5-Fluorouracil, on DNMT1 from Decitabine, and on TOP1 from Irinotecan. All five appear as direct effects at 24 or 48 hours and none at 6 hours, which fits mechanisms that need enzyme turnover or DNA replication. Four further pairs show the expected direction in the direct effects without reaching significance after correction. TYMS increases after Pemetrexed at 24 and 48 hours (uncorrected $p = 5.5 \times 10^{-5}$ and $4.4 \times 10^{-4}$), FKBP5 increases after Hydrocortisone and after Medroxyprogesterone Acetate at 24 and 48 hours (uncorrected $p$ between $2.1 \times 10^{-4}$ and 0.05), and Azacytidine lowers DNMT1 at 24 and 48 hours (uncorrected $p = 0.006$

and 0.035), less than Decitabine, as expected from its longer route into DNA. TOP2B decreases after Doxorubicin and Epirubicin at 48 hours only in the total effects (uncorrected $p = 0.03$ and 0.006), and POLR2A shows no significant decrease after Lurbinectedin at any time point. None of the nine pairs predicted to stay unchanged is significant at 24 or 48 hours. This holds for the five EGFR inhibitors on EGFR, Capecitabine on TYMS, and the three PARP inhibitors on PARP1, consistent with the prediction, although a non-significant test does not exclude a small effect. At 6 hours, PARP1 increases after Niraparib and Talazoparib, but also after ten other drugs, among them two CDK4/6 inhibitors, two aromatase inhibitors, and an alkylating agent, so this early increase might belong to the shared 6-hour response rather than to PARP trapping.

The drug pairs show one result with a known mechanism. Cisplatin and 5-Fluorouracil raise TYMS at 24 hours beyond the sum of their single effects ($p = 6.0 \times 10^{-6}$). Cisplatin enlarges the pool of reduced folates, which increases the formation of the complex between the 5-Fluorouracil metabolite and thymidylate synthase [25], and more inhibitor-bound enzyme is what the two mechanisms above turn into more protein. The other 23 significant interactions on the set are at 6 hours and negative, 15 of them on POLR2A, so the shared early response to a drug pair is smaller than the sum of the responses to its two drugs.

The temporal network adds little for this set. At a false discovery rate of 5%, seven of the nine proteins depend on their own earlier expression, most strongly TYMS from 24 to 48 hours ($p = 3 \times 10^{-35}$), and none of the 40 remaining edges links two of the nine proteins. TOP1 at 24 hours precedes six proteins at 48 hours with positive coefficients, among them RPA1 and PAICS, proteins of DNA replication and purine synthesis ($p = 0.002$ and 0.013), which suggests that TOP1 abundance follows the replicative state of the cells.

For exploration, the level can be relaxed, as the main text discusses for the uncorrected p-values. At a false discovery rate of 15%, PARP1 at 6 hours precedes 15 proteins at 24 hours, all with positive coefficients, and six of them belong to RNA processing, HNRNPD, SNRPB, SARNP, SMU1, EFTUD2, and CPSF6, which is consistent with the identification of heterogeneous nuclear ribonucleoproteins as poly(ADP-ribose)-binding proteins [26]. Of the six, only the edge to HNRNPD is significant at 5%, and we list the group as a hypothesis, not as a finding.

Taken together, `Drug-Prot` recovers five of the twelve predicted changes, for four target enzymes, with the direction, time course, and specificity that the mechanisms predict, and none of the nine pairs predicted to stay unchanged shows a significant change at 24 or 48 hours. It misses the predicted degradation of TOP2B and POLR2A, which may be transient at the measured time points or diluted over the 18 cell lines.
